# Transgenerational effects decrease larval resilience to ocean acidification & warming but juvenile European sea bass could benefit from higher temperatures in the NE Atlantic

**DOI:** 10.1101/2021.11.15.468704

**Authors:** Sarah Howald, Marta Moyano, Amélie Crespel, Louise Cominassi, Guy Claireaux, Myron A. Peck, Felix C. Mark

**Affiliations:** Alfred Wegener Institute Helmholtz Centre for Polar and Marine Research, Integrative Ecophysiology, Bremerhaven, Germany; Institute of Marine Ecosystem and Fisheries Science, Center for Earth System Research and Sustainability (CEN), University of Hamburg, Germany; Center for Coastal Research, University of Agder, Postbox 422, 4604 Kristiansand, Norway; Université de Bretagne Occidentale, LEMAR (UMR 6539), Brest, France; Ifremer, LEMAR (UMR 6539), Laboratory of Adaptation, and Nutrition of Fish, Centre Ifremer de Bretagne, Plouzané, France; Coastal Systems (COS), Royal Netherlands Institute for Sea Research (NIOZ), Netherlands; Institute of Arctic Biology, University of Alaska, Fairbanks, PO Box 757000, Fairbanks, AK 99775, USA

**Keywords:** Dicentrarchus labrax, ocean acidification, ocean warming, metabolic rates, larval growth, juvenile growth, teleost

## Abstract

1.

The aim of this study was to investigate the effect of ocean acidification (OA) and warming (OW) as well as the transgenerational effect of OA on larval and juvenile growth and metabolism of a large economically important fish species with a long generation time. Therefore we incubated European sea bass from Brittany (France) for two generations (>5 years in total) under current and predicted OA conditions (*P*CO_2_: 650 and 1700 µatm). In the F1 generation both OA condition were crossed with OW (temperature: 15-18°C and 20-23°C). We found that OA alone did not affect larval or juvenile growth and OW increased developmental time and growth rates, but OAW decreased larval size at metamorphosis. Larval routine metabolic rate (RMR) and juvenile standard metabolic rate (SMR) were significantly lower in cold compared to warm conditioned fish and also lower in F0 compared to F1 fish. We did not find any effect of OA on RMR or SMR. Juvenile *P*O_2crit_ was not affected by OA, OW or OAW in both generations.

We discuss the potential underlying mechanisms resulting in beneficial effects of OW on F1 larval growth and RMR and in resilience of F0 and F1 larvae and juveniles to OA, but on the other hand resulting in vulnerability of F1, but not F0 larvae to OAW.. With regard to the ecological perspective, we conclude that recruitment of larvae and early juveniles to nursery areas might decrease under OAW conditions but individuals reaching juvenile phase might benefit from increased performance at higher temperatures.

**Summary statement:** We found that OA did not affect developmental time, growth, RMR and SMR, while OW increased these traits. OAW decreased larval size at metamorphosis. We discuss underlying mechanisms and the ecological perspective resulting from these results and conclude that recruitment to nursery areas might decrease under OAW conditions but individuals reaching juvenile phase might benefit from increased performance at higher temperatures in Atlantic waters.

## 3. Introduction

Climate change is leading to increasing ocean surface temperatures (ocean warming – OW), as well as decreasing ocean pH (ocean acidification – OA). OW as a single stressor on fish metabolism has been investigated intensively since the 1980s in a variety of fish species and life stages and directly influences their metabolism and therefore their growth (Johnson & Katavic, 1986; Peck, 2002; Pörtner, et al., 2007), reproduction success (see review Llopiz, et al., 2014), as well as distribution range and abundance (Turner, et al., 2009; Pörtner, 2006). OW can increase growth rates of larval and juvenile fish (Chauton, et al., 2015; Baumann, 2019), within their thermal window. Although studies on larvae are less numerous than those on adults and juveniles, it has become obvious that larvae are less resilient to OW than adults and juveniles (Dahlke, et al., 2020).

Initially, fish had been thought to be less vulnerable to OA due to well-developed acid-base regulation systems (Heuer & Grosell, 2014), yet their capacity to cope with OA and ocean acidification and warming (OAW) as co-occurring stressors has been investigated intensively during the last decade with species and life stage specific results (Cattano, et al., 2017): OA levels between 700 and 1600 µatm CO_2_ can lead to increased larval growth (mahi-mahi, Bignami, et al., 2014; clownfish, Munday, et al., 2009), but decreased larval swimming performance (mahi-mahi, Bignami, et al., 2014; dolphinfish, Pimentel, et al., 2014) and larval metabolic rates (dolphinfish, Pimentel, et al., 2014). OA also induced severe to lethal tissue damage (cod larvae, Frommel, et al., 2011) and increased larval otolith size, with possible implications for hearing sensitivity (cobia and mahi-mahi, Bignami, et al., 2013; 2014, respectively). In other species growth was decreased by OA (inland silverside juveniles, Baumann, et al., 2012), or not affected (Atlantic halibut juveniles, Gräns, et al., 2014; cobia larvae, Bignami, et al., 2013). Dahlke et al. (2020) showed that Atlantic cod embryos demonstrated poor acid base regulation capacities before and during gastrulation, connected to increased mortality under OA and OAW. On the contrary, acid base regulation capacities after gastrula were similar to that of adult cod. If both stressors were combined, the effects became more unidirectional and were synergistic in most fish species, e.g. OAW increased growth and survival in larval and juvenile sea bass in their Atlantic populations, but decreased physiological performance (Pope, et al., 2014). The cumulative consequences of these changes are to be determined.

An important factor for projecting whether a species will be able to keep their distribution range under changing conditions, is their potential and capacity to acclimate and adapt over generations. Few studies have so far examined transgenerational effects of fish in the context of OAW, with trait- and species-specific capacities to adapt to future conditions. For example, in cinnamon anemone fish (*Amphiprion melanopus*) the negative effect of OA on escape responses was reduced in some traits if parents were exposed to OA (Allan, et al., 2014), whereas in spiny damselfish (*Acanthochromis polyacanthus*), negative effects on olfactory responses were not reduced after parental exposure to OA (Welch, et al., 2014). In addition to the low number of studies on transgenerational effects, they usually used small fish, with short generation times and applied only one stressor, either OW or OA. Little is known about the combined effect of several stressors on economically important larger-sized fish with longer generation times and thus multi-stressor, transgenerational studies on such fish are necessary to project future distribution of fish.

Consequently, in our study we used European sea bass *Dicentrarchus labrax* as a larger, long-lived model species. Sea bass is an economically important species in industrial and recreational fishing as well as in aquaculture (160 000 t in 2015, Bjørndal & Guillen, 2018). Generally rather resilient towards environmental fluctuations, effects of OW and OA have been reported for several seabass life stages: OW increased growth rates in larval sea bass, although at the expense of decreased swimming performance (Atlantic population, 15 to 20°C, Cominassi, et al., 2019). Exposure to OA throughout larval development increased mineralization and reduced skeletal deformities (Atlantic population, 19°C and 15 and 20°C, respectively, Crespel, et al., 2017; Cominassi, et al., 2019). In combination, OAW did not have additional effects on larval growth, swimming ability and development than those already observed separately (Atlantic population, Cominassi, et al., 2019). Juvenile sea bass are highly tolerant to temperature (Dalla Via, et al., 1998; Claireaux & Lagardère, 1999) and show some degree of tolerance to OA as a single stressor at the mitochondrial level (Atlantic population, Howald, et al., 2019). OA and OW acted antagonistically: OW as a single stressor increased growth and digestive efficiency, while OA did not affect these traits. Both stressors combined resulted in reduced growth and digestive efficiency compared to the impact of OW alone. Low food ratios enhanced this effect resulting in an even more pronounced growth and digestive efficiency reduction than under OAW alone (Atlantic population, Cominassi, et al., 2020).

This study aimed to investigate the effect of OAW as well as the transgenerational effect of OA on larval and juvenile growth and metabolism. Therefore, we incubated sea bass from an Atlantic population for two generations (>5 years in total) under current and predicted OA conditions (*P*CO_2_: 650 and 1700 µatm) and applied a warming condition on larvae and juveniles of the F1 generation (ambient, 15-18°C, and Δ5°C, 20-23°C). To study the effect of OW (F1), OA (F0, F1) and OAW (F1) on sea bass, we investigated growth (F0, F1) through ontogeny as a proxy for whole organism fitness. In addition we measured routine metabolic rates (RMR, F1) of larvae, as well as standard metabolic rates (SMR, F0, F1) and critical oxygen concentration (*P*O_2crit_, F0, F1) of juvenile sea bass, to unravel the underlying mechanisms resulting in possible growth differences. In F0, no effect of OA on larval and juvenile growth or juvenile SMR and *P*O_2crit_ were found (Crespel, et al., 2017; 2019). Those traits were compared in F0 and F1 fish to determine transgenerational effects due to parental acclimation to different OA levels. Our hypotheses were: (1) OW will lead to increased growth and metabolic rates in F1 larvae and juveniles. (2) OA alone will not have significant effects on larval and juvenile growth and metabolism in F1, as sea bass seem to be quite tolerant to OA and no detrimental effects were found in F0. (3) In combination, OA will lead to synergistic OAW effects, reflected in lower growth in larvae and juveniles.

## 4. Materials and Methods

The present work was performed within the facilities of the Ifremer-Centre de Bretagne (agreement number: B29-212-05). Experiments were conducted according to the ethics and guidelines of the French law and legislated by the local ethics committee (Comité d’Ethique Finistérien en Experimentation Animal, CEFEA, registering code C2EA-74) (Authorizations APAFIS 4341.03, #201620211505680.V3 and APAFIS 14203-2018032209421223 for F0 and F1, respectively).

### 4.1. Animals and experimental conditions

Sea bass were reared from early larval stage onwards in two OA treatments in F0 and four OAW treatments in F1. F0 fish were reared in two OA scenarios, following the predictions of the Intergovernmental Panel on Climate Change (IPCC, 2021) for the next 130 years: today’s ambient situation in coastal waters of Brittany and the Bay of Brest (A, approx. 650 µatm (cf. Pope, et al., 2014; Duteil, et al., 2016)) and a scenario according to SSP5-8.5, projecting a Δ*P*CO_2_ of 1000 µatm (Δ1000, approx. 1700 µatm). Adults from these two treatments were used in the reproduction experiments to generate F1. Sea bass of F1 were reared under the same OA conditions as their respective parents. Additionally two different temperatures were applied on each OA condition in F1 to create a cold and a warm life condition scenario or four OAW conditions (C-A, C-Δ1000, W-A and W-Δ1000), respectively.

Larval rearing was performed in a temperature controlled room and water temperatures were fixed to 19°C in F0, and 15 and 20°C in F1-C and F1-W, respectively. In juveniles and adults, water temperatures of F0 and F1-C sea bass were adjusted to ambient temperature in the Bay of Brest during summer (up to 19°C), but were kept constant at 15 and 12°C for juveniles and adults, respectively, when ambient temperature decreased below these values. F1-W was always 5°C warmer than the F1-C treatment. A flow chart summarizing temperature and *P*CO_2_ conditions as well as replicate tank number, tank volume and number of individuals per tank is shown in Figure 1.

**Figure 1.**
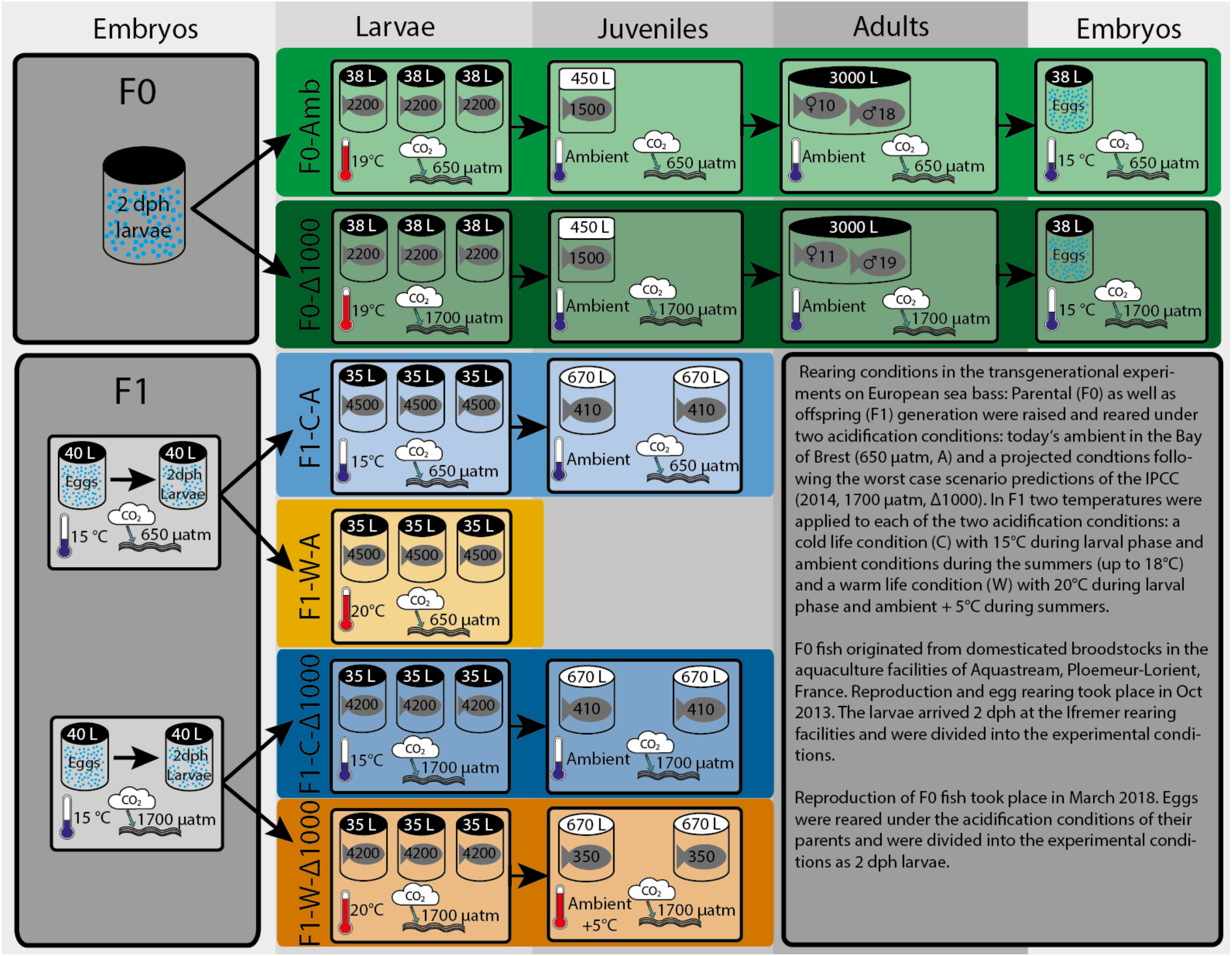
Schematic overview summarizing the rearing conditions of two generations of sea bass under different OAW scenarios.

During larval rearing, the photoperiod was set to 24h darkness during the first week and 16h light and 8h darkness (12h each in F1, respectively) per day afterwards. Light intensity increased progressively during the larval rearing period from total darkness to about 100 lux (Table S 1). To work in the larval rearing facilities, headlamps were used (set to lowest light intensity). In the juvenile and adult rearing facilities photoperiod followed natural conditions (adjustment once a week).

#### 4.1.1. F0 generation

##### 4.1.1.1. Larval rearing

F0 Larval rearing and origin is described in detail in Crespel et al. (2017; 2019), briefly, larvae were obtained from the aquaculture facility Aquastream (Ploemeur-Lorient, France) at 2 dph (October 2013). Brood stock derived from a population selected for growth for five generations. Three blocks of four females (mean weight 5.6 kg) were crossed reciprocally with five to seven males (mean weight 3.7 kg). Conditions in the aquaculture facility during egg rearing were as follows: 14±0.2°C, 38 psu, pH 7.9±0.1. After transferring to the Ifremer facilities, larvae were randomly distributed among the two OA conditions described above. Larvae were reared in nine black 38 L tanks initially stocked with *ca*. 2200 larvae tank^-1^ in triplicates for all conditions. Larvae were fed *ad libitum* via continuous delivery of Artemia nauplii until 28 dph. Afterwards, commercial dry pellets (Neo Start, LeGouessant, France) were fed for the rest of the larval period.

##### 4.1.1.2. Juvenile rearing

Juvenile rearing was described in detail in Crespel et al. (2019). As larvae and post-larval juveniles would display different growth rates under the different life condition scenarios, we adopted the concept of degree days (dph · T(°C)) as a basis for comparison between them. Briefly, the early juveniles were counted per tank and transferred from larval to juvenile rearing facilities at approx. 820 degree-days (dd) (45 dph). Juveniles of one condition were combined and kept in square shaped 450L tanks (n=1500 fish per condition). At 8 months (about 250 dph), juveniles were PIT tagged (marked with passive integrated transponders). Juveniles were fed daily with commercial fish food (Neo Start), which was adjusted in size and amount, as recommended by the supplier (Le Gouessant, Lamballe, France). Food ratios were adjusted after each sampling for growth, approx. every 30 days or 3-4 weeks in F0 and F1, respectively (see below), using the formulae provided by Le Gouessant. Daily food ratios were supplied to the tanks by automatic feeders during day time.

##### 4.1.1.3. Broodstock rearing

During the reproductive season 2017 (fish were 3.5 years old), sex steroid plasma concentration was measured regularly in all broodstock fish. The individuals with the highest concentrations were kept in round black tanks with a volume of 3 m³ and a depth of 1.3 m. Each of the two tanks (one for each condition) was stocked with 22 males and 11 females, resulting in fish density of 11.6 kg m^-3^ and 11.0 kg m^-3^ in A and Δ1000, respectively. Mass and length were regularly measured and commercial fish food was adjusted accordingly. Fish were fed Vitalis CAL (Skretting, Norway) during reproduction season and Vitalis REPRO (Skretting, Norway) during the rest of the year. Vitalis REPRO was supplied to the tanks with automatic feeders during daytime. Vitalis CAL was supplied to the tank manually in three to four rations during week days.

#### 4.1.2. F1 generation

Embryos were obtained by artificial reproduction of F0 broodstock fish. Succinctly, once the water temperature reached 13°C and the first naturally spawned eggs were observed in the egg collectors, females were injected with gonadotropin-releasing hormone (GnRH, 10 µg kg^-1^) to accelerate oocyte maturation (23.03.2018). After three days (26.03.2018) eggs and milt were stripped from ripe females and males, respectively, and artificial fertilization was performed following the protocol of Parazo et al. (1998). Briefly, eggs (10 ml l^-1^) were mixed with sea water and milt (0.05 ml milt L^-1^ seawater). Ten females (1.56 ± 0.24 kg) were crossed with 18 males (1.07 ± 0.16 kg) and 11 females (1.28 ± 0.30 kg) were crossed with 19 males (0.99 ± 0.19 kg) in the A and Δ1000 groups, respectively. Fertilized eggs were incubated in 40 L tanks (without replicates) at 15°C and at the same *P*CO_2_ conditions as respective F0. Hatching occurred after four days (30.03.2018).

##### 4.1.2.1. Larval rearing

Two days after hatch (02.04.2018), larvae were distributed into twelve black 35 L tanks. Triplicate tanks were allocated to each of the four OAW treatments with *ca*. 4500 and 4200 larvae tank^-1^ in A and Δ1000 tanks, resulting in a total of *ca.* 13500 and 12800 larvae condition^-1^ in A and Δ1000, respectively. The temperature of the tanks allocated to warm life condition was increased stepwise during the following five days, by 1 °C day^-1^. Starting at 7 days post-hatch (dph) (mouth opening), larvae were fed with live artemia, hatched from High HUFA Premium artemia cysts (Catvis, AE ‘s-Hertogenbosch, Netherlands). Artemia were fed to the larvae 24h after rearing cysts in sea water. Larvae were fed *ad libitum* with artemia during the day, excess artemia left the tank via the waste water outflow. Larval mortality was 26-96 %, without any pattern for OAW condition (Table S 2). High mortality of sea bass larvae, especially during early larval rearing are common in science and aquaculture (e.g. Nolting, et al., 1999; Suzer, et al., 2007; Villamizar, et al., 2009). We could not find any signs of infection neither in the tanks with high mortality, nor in the tanks with lower mortality rates. However, as larval mortality was unreasonably high (96%) within the first week in one of the replicate tanks of the W-A treatment, remaining larvae in this tank were euthanized (sedation followed by an anaesthetic overdose) and not used for further analysis. Water surface was kept free of oily films using a protein skimmer. Water exchange was set to 25 l/h and stepwise increased to 40 l/h at the end of larval rearing.

##### 4.1.2.2. Juvenile rearing

At approx. 950 dd, the early juveniles were counted per tank and transferred from larval to juvenile rearing (48 dph, 17.05.2018 and 63 dph, 01.06.2018 for W and C, respectively). For F1-W, only the Δ1000 fish were transferred to juvenile rearing facilities. Juveniles were randomly allocated to duplicate tanks per condition. Swim bladder test was done at 1680 dd (83 dph, 21.06.2018) and 1661 dd (104 dph, 12.07.2018) for F1-W and F1-C, respectively. Briefly, the fish were anaesthetized and introduced into a test container with a salinity of 65 psu (Marine SeaSalt, Tetra, Melle, Germany). In F1-W, all floating fish with a developed swim bladder were counted and kept in the rearing tanks, resulting in 355 fish per tank (710 fish in total). In F1-C, 410 fish per tank were randomly selected (820 fish per condition), to have similar stocking densities in W and C. Non-floating fish as well as excess F1-C fish were counted and euthanized (sedation followed by an anesthetic overdose). The juveniles were reared in round tanks with a volume of 0.67 m^3^ and a depth of 0.65 m. During the first five days after moving to juvenile rearing, the juveniles were fed artemia nauplii and commercial fish food. Afterwards commercial fish food was fed as described above.

#### 4.1.3. Experimental conditions

##### 4.1.3.1. Sea water preparation

The sea water used in the aquaria was pumped in from the Bay of Brest from a depth of 20 m approximately 500 m from the coastline, passed through a sand filter (∼500 µm), heated (tungsten, Plate Heat Exchanger, Vicarb, Sweden), degassed using a column packed with plastic rings, filtered using a 2 µm membrane and finally UV sterilized (PZ50, 75W, Ocene, France) assuring high water quality.

Water conditions for the rearing tanks were preadjusted to the desired OAW condition in header tanks. Sea water arrived in a reservoir next to the rearing facilities, after passing the tungsten heater, in F1, two different reservoirs were used, to create the different temperature conditions. The temperature controlled water supplied the header tanks within the rearing facilities to adjust the water to the desired OA condition. Each header tank supplied water to all replicate tanks of the respective condition.

In F0 larvae and juveniles the water pH in the header tank was controlled by an automatic injection system connected to a pH electrode (pH Control, JBL, Germany), which injected either air (A) or CO_2_ (Δ1000), to control water pH. For the Δ1000 F1 larvae the CO_2_-bubbling was installed in the middle of the header tank and the water was mixed continuously with a pump. The CO_2_-bubbling was adjusted by a flow control unit, when pH deviated from the desired value.

Older F0-A juveniles (> 2 years) and adults, as well as F1-A larvae and juveniles received water directly from the respective reservoir, without header tank. Additionally as water exchange rates became too high for the automatic injection system and the header tank, PVC columns were installed to control the pH in the rearing tanks. The temperature controlled water arrived at the top of the column and was pumped from the bottom of the column to the rearing tanks. The CO_2_-bubbling was installed at the bottom of the column and was adjusted by a flow control unit, when pH deviated from the desired value.

##### 4.1.3.2. Calculation of water chemistry

The Microsoft Excel macro CO2sys (Lewis & Wallace, 1998) was used to calculate seawater carbonate chemistry, the constants after Mehrbach et al. (1973, as cited in CO2sys) refit by Dickson and Millero (1987, as cited in CO2sys), were employed.

From October 2015 onwards (late juveniles of F0), total alkalinity was measured following the protocol of Anderson & Robinson (1946) and Strickland & Parsons (1972): 50 ml of filtered tank water (200 µm nylon mesh) were mixed with 15 ml HCl (0.01 M) and pH was measured immediately. Total alkalinity was then calculated with the following formula:

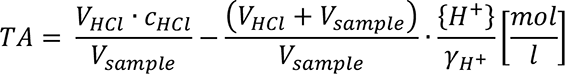

With: TA – total alkalinity [mol · l^-1^], V_HCl_ – volume HCl [l], c_HCl_ – concentration HCl [mol · l^-1^], V_sample_ – volume of sample [l], H^+^ – hydrogen activity (10^-pH^), γ^H+^ – hydrogen activity coefficient (here γ^H+^ = 0.758).

##### 4.1.3.3. Water quality control

Temperature and pH were checked each morning with a handheld WTW 330i or 3110 pH meter (Xylem Analytics Germany, Weilheim, Germany; with electrode: WTW Sentix 41, NIST scale) before feeding the fish. Until F0 juveniles reached 2 years, the pH meter and the automatic injection system were calibrated weekly with fresh buffers (Merk, Germany). Measured values never differed more than 2% from the target values. Afterwards the pH meter was calibrated daily with NIST certified WTW technical buffers pH 4.01 and pH 7.00 (Xylem Analytics Germany, Weilheim, Germany).

Total pH was determined twice during F0 larval rearing (start and end) and nine times during F0 juvenile rearing following Dickson et al. (2007) using m-cresol purple as indicator. Additionally, water samples were sent to LABOCEA (France) to measure total alkalinity by titration, as well as phosphate and silicate concentration by segmented flow analysis following Aminot et al. (2009).

In later F0 juveniles (> 2 years) and adults as well as F1 larvae and juveniles, total alkalinity was measured monthly or weekly in F0 and F1, respectively, following the protocol described above. Oxygen saturation (WTW Oxi 340, Xylem Analytics Germany, Weilheim, Germany) and salinity (WTW LF325, Xylem Analytics Germany, Weilheim, Germany) were measured together with total alkalinity (monthly in F0 and weekly in F1). The tanks were cleaned daily after pH-measurements. Water flow within the tanks was adjusted once a week, so that oxygen saturation levels were kept >85%, with equal flow rates in all tanks of one temperature. All water parameters are summarized in Table 1 for F0 larvae and juveniles and Table 2 for F0 broodstock (two years before spawning) and F1 larvae and juveniles.

**Table 1.** Water parameters during larval and early juvenile phase of F0: Larval period until (45 dph, ∼900 dd), early juveniles until 1.5 years. Means ± s.e.m. over all measurements per condition (triplicate tanks in larvae, single tanks in juveniles). Temperature (Temp.) and pH (NBS scale) were measured daily. pH (total scale), salinity, phosphate, silicate and total alkalinity (TA) were measured once at the beginning and once at the end of the larval phase and 9 times during juvenile phase; *P*CO_2_was calculated with CO2sys; A–Ambient *P*CO_2_, Δ1000 –ambient + 1000 µatm CO_2_,L – Larvae, J – Juveniles, (see Crespel, et al., 2017; Crespel, et al., 2019).

| Treatment | pH<sub>NBS</sub> [-] | pH<sub>total</sub> [-] | Temp.<br>[°C] | Salinity<br>[psu] | TA<br>[μmol L<sup>-1</sup>] | $PCO_2$<br>[μatm] | $PO_4^{3-}$<br>[μmol L<sup>-1</sup>] | $SiO_4$<br>[μmol L<sup>-1</sup>] |
| --- | --- | --- | --- | --- | --- | --- | --- | --- |
| L A | 7.96±0.01 | 7.89±0.01 | 19.2±0.3 | 33.8±0.2 | 2294±3 | 589±10 | 0.57±0.01 | 8.94±0.06 |
| L $\Delta 1000$ | 7.59±0.00 | 7.54±0.03 | 19.2±0.3 | 33.8±0.2 | 2306±9 | 1521±97 | 0.57±0.01 | 8.94±0.06 |
| J A | 8.05±0.01 | 7.94±0.03 | 15.3±0.1 | 34.3±0.2 | 2294±10 | 516±31 | 0.71±0.08 | 8.35±0.26 |
| J $\Delta 1000$ | 7.61±0.01 | 7.53±0.02 | 15.3±0.1 | 34.3±0.2 | 2280±16 | 1489±42 | 0.71±0.08 | 8.35±0.26 |

**Table 2.** Water parameters in the 2 years before spawning of F0 (2016-2018) and during larval and juvenile phase of F1: Larval period until 17.05.2018 (48 dph, ∼900 dd) and 01.06.2018 (63 dph, ∼900 dd) for warm and cold life condition respectively, for the juveniles until 28.09.2018 (180 dph, ∼4000 dd) and 12.02.2019 (319 dph, ∼5100 dd) for warm and cold conditioned fish respectively. Means ± s.e. over all replicate tanks per condition. Temperature (Temp.), pH (free scale), salinity, oxygen and total alkalinity (TA) were measured weekly in F1 and monthly in F0; *P*CO_2_ was calculated with CO2sys; sea water (SW) measurements were conducted in 2017 and 2018; A – Ambient *P*CO_2_, Δ1000 – ambient + 1000 µatm CO_2_, L – Larvae, J – Juveniles, C – cold life condition, W – warm life condition.

| Treatment | pH<sub>free</sub><br>[-] | Temp.<br>[°C] | Salinity<br>[psu] | O<sub>2</sub><br>[% airsat.] | TA<br>[-] | PCO<sub>2</sub><br>[$\mu\text{atm}$] |
| --- | --- | --- | --- | --- | --- | --- |
| F0 A | 7.95±0.02 | 14.1±0.6 | 33.6±0.3 | 92.4±1.7 | 2406±49 | 670±40 |
| F0 $\Delta 1000$ | 7.59±0.02 | 14.1±0.6 | 33.6±0.3 | 92.4±1.9 | 2411±46 | 1616±74 |
| F1 L C A | 8.06±0.01 | 15.3±0.1 | 31.8±0.1 | 94.3±1.0 | 2360±23 | 504±19 |
| F1 L C $\Delta 1000$ | 7.53±0.01 | 15.5±0.1 | 31.8±0.1 | 94.3±0.8 | 2330±22 | 1872±74 |
| F1 L W A | 7.96±0.01 | 20.2±0.2 | 31.7±0.0 | 84.9±3.4 | 2311±32 | 656±22 |
| F1 L W $\Delta 1000$ | 7.61±0.01 | 20.2±0.2 | 31.8±0.0 | 88.1±1.7 | 2321±32 | 1624±59 |
| F1 J C A | 7.94±0.01 | 16.1±0.2 | 33.0±0.1 | 92.4±0.5 | 2376±15 | 696±19 |
| F1 J C $\Delta 1000$ | 7.60±0.01 | 16.3±0.2 | 33.0±0.1 | 94.3±0.5 | 2380±14 | 1603±32 |
| F1 J W A | n.a. | n.a. | n.a. | n.a. | n.a. | n.a. |
| F1 J W $\Delta 1000$ | 7.57±0.02 | 22.7±0.2 | 33.0±0.2 | 86.3±1.3 | 2323±16 | 1866±83 |
| SW | 8.07±0.01 | 15.0±0.5 | 34.6±0.3 | 101.0±0.8 | 2441±23 | 609±37 |

### 4.2. Growth

#### 4.2.1. Larval growth

##### 4.2.1.1. F0 larvae

Larval growth was measured as described in Crespel et al. (2017). Briefly, 10 larvae per tank were sampled each week, starting at 15 dph and ending at 45 dph, when 30 larvae per tank were sampled. For growth measurements, larvae were anaesthetized with phenoxyethanol (200 ppm) and their wet mass (WM), as well as body length (BL) were measured. BL in F0 larvae was measured with a caliper from the tip of the snout to the end of the notochord until flexion, afterwards fork length was considered as BL, see Figure S 1.

##### 4.2.1.2. F1 larvae

In F1 larvae, individuals were sampled every 200 dd from 100 – 900 dd to follow growth throughout the larval phase. At each sampling point, 20 larvae per tank were anaesthetized with MS-222 (50 mg l^-1^, Pharma Q) prior to feeding and directly photographed individually with a microscope (Leica M165C). The larvae were then frozen in liquid nitrogen and stored at −80°C until dry mass (DM) measurements. The software ImageJ (Schneider, et al., 2012) was used to determine BL of larvae, see Figure S 1 on the definition of BL.

#### 4.2.2. Growth in juveniles

BL and WM were measured approx. every 30 days in F0 and every 3 – 4 weeks in F1 juveniles. Early juveniles were starved for one day prior to growth samplings. Later on, two days of starving were put into practice, to make sure that digestive tracts were empty. Juveniles were caught from their tanks and anaesthetized with MS-222 (Pharma Q). Concentration of anesthetic was adjusted to reach a loss of equilibrium within less than 5 minutes, typically 0.2 g l^-1^. WM and BL were directly determined with a precision balance (Sartorius MC1 AC210P) and calipers. For all sampling, only the morning hours were used, to avoid diurnal artefacts in data.

#### 4.2.3. Data handling

For F1 larvae and juveniles, mean specific growth rates (SGR [% day^-1^]) of each tank were calculated after Sutcliffe (1970) with the following formula:

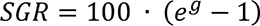

The instantaneous growth coefficient (g) was calculated as followed:

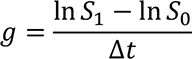

With: S_0_ and S_1_ – initial and final size (BL, WM or DM) and Δt – time between the two measurements [days]. Initial and final sizes were calculated for three quantiles (0.05, 0.5 and 0.95) for each tank (“ecdf” function in R).

Q_10_ was calculated with the following formula:

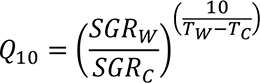

With: SGR – specific growth rate, T – temperature, W and C as subscripts for W and C condition.

### 4.3. Respirometry

#### 4.3.1. F1 larvae

Larval respiration measurements were conducted from approx. 350 to 950 dd in all conditions (18 – 47 dph and 25 – 63 dph in W and C, respectively).

Larval respiration was measured in an intermittent flow system. The setup consisted of up to eight 4 ml micro respiration chambers with a glass ring (Unisense A/S, Aarhus, Denmark), equipped with a glass coated magnetic stirrer (Loligo® Systems, Viborg, Denmark) and a stainless steel mesh (Loligo® Systems, Viborg, Denmark), to separate the stirrer from the larva. The magnetic stirrers were connected to one stirrer controller (Rank Brothers Ltd., Cambridge, England). The chamber was closed with a custom-made glass lid with three metal ports: two with a diameter of 0.8 mm for water inflow and outflow during the flushing, and one with 1.2 mm to insert the oxygen sensor into the chamber. Oxygen concentration within the chamber was measured with oxygen microsensors connected to a FireSting oxygen meter (PyroScience GmbH, Aachen, Germany). The respiration chambers were placed within a rack without shielding between the individual chambers. The rack holding the respiration chambers was fully submerged in a water reservoir, which received flow through water from the respective header tanks of the larval rearing. Water conditions within the water reservoir were kept at 15.5±1.5 °C and 21.2±1.0°C for W and C larvae, combined with the OA condition of the origin tank of the respective larvae. The reservoir was a black container, which shielded the respiration setup from external disturbances. During the flushing periods, water from the reservoir was pumped into the respiration chambers using computer-controlled flush pumps (Miniature DC pump, Loligo® Systems, Viborg, Denmark), relays and software (AquaResp, Copenhagen, Denmark). Four chambers were connected to one flush pump and controlled by one computer. Oxygen microsensors were calibrated to 0% saturation (nitrogen purged seawater) and 100% saturation (fully aerated seawater) prior to each measurement.

Respiration measurements were done in the larval rearing facilities with the same light conditions as for larval rearing. Larvae were fasted at least three hours prior to respiration measurements to minimize the effect of specific dynamic action (SDA) on metabolic rate. Preliminary tests with measurements overnight proved that oxygen consumption during the 12 h after the 3-h fasting period was similar, suggesting no contribution of SDA and thus that the 3h-fasting was sufficient for our setup. Larvae were individually placed in the respiration chambers. Oxygen partial pressure was measured every second for approx. four hours. Cycles were composed of 420 s flush, followed by 60 s wait time (time after flush pump stopped to wait for stable drop in oxygen concentration) and 600 to 180 s measurement time (13-20 cycles per larvae). Measurement time was decreased with increasing larval size. Oxygen concentration was restored to normoxia during the flush time of each cycle and was usually kept above 75% air saturation. Background respiration was measured for 30 min (one slope) after 11 and 18 measurements in F1-C and F1-W larvae, respectively. The mean bacterial respiration was calculated for each temperature treatment and subtracted from total respiration of all larvae of this temperature to obtain oxygen consumption of the larva. Background respiration was typically 0.5 - 6 % of total respiration. Only declines in oxygen concentration displaying R² > 0.80 were used for analysis. After the measurement, larvae were checked if alive, anaesthetized with MS-222 (50 mg l^-1^ Pharma Q), photographed individually and frozen in liquid nitrogen. Length and DM of the larvae was obtained as described above (see Table S 3). After each experiment, the respiration system was rinsed with fresh water and let dry. For disinfection, respiration chambers, the tubing of the flush pump and the oxygen sensors were additionally rinsed with ethanol, which was allowed to sit in the chambers and the tubing for at least 30 min followed by rinsing with distilled water.

#### 4.3.2. Juveniles

##### 4.3.2.1. Set up F0 juveniles

Measurements on the 15 months old F0 juveniles (F0-old) were described in Crespel et al. (2019), measurements on the 5 months old F0 juveniles (F0-young) were done similarly and if different, the information for F0-young are given in brackets. Briefly, F0 juvenile respiration was measured individually in one of four (eight) intermittent flow respirometry chambers with a volume of 2.1 l (60 ml), which were submerged in a tank which received flow-through seawater at 15 ± 0.25 °C and the respective acidification condition. The water was recirculated within the chamber with a peristaltic pump with gas-tight tubing. The oxygen probe (FireSting oxygen meter, PyroScience GmbH, Aachen, Germany or multichannel oxygen meter, PreSens Precision Sensing GmbH, Regensburg, Germany) was placed within the recirculation loop. Oxygen sensors were calibrated to 0% saturation (sodium sulfite, saturated) and 100% saturation (fully aerated seawater) prior to each experiment. The flush pumps were controlled by relays and software (AquaResp, Copenhagen, Denmark). The setup was placed behind a curtain to avoid disturbances. Background respiration was measured after each experiment and estimated for the whole experiment by linear regression assuming zero background respiration at the beginning of the run as the entire system was disinfected with household bleach between each trial.

##### 4.3.2.2. Set up F1 juveniles

F1 juvenile respiration was measured in an intermittent flow system. The setup consisted of up to eight 450 ml custom-made respiration chambers. The chambers were made from Lock&Lock glass containers with plastic lid. Four rubber ports were placed into the lid: two for water inflow and outflow during flushing cycles and two to connect the chamber to a mixing pump (Miniature DC pump, Loligo® Systems, Viborg, Denmark). Oxygen concentration was measured with robust oxygen probes placed within the circulation loop and connected to a FireSting oxygen meter (PyroScience GmbH, Aachen, Germany) or to a multichannel oxygen meter (PreSens Precision Sensing GmbH, Regensburg, Germany). The respiration chambers were fully submerged in a flow-through water reservoir. Water conditions within the water reservoir were kept at 14.9±1.0 °C and 22.3±1.8°C for C and W larvae, combined with the OA condition of the origin tank of the respective juvenile. During the flushing periods, water from the reservoir was pumped into the respiration chambers using computer-controlled flush pumps (EHEIM GmbH & Co. KG, Deizisau, Germany), relays and software (AquaResp, Copenhagen, Denmark). Four chambers were connected to one flush pump and controlled by one computer, running either the FireSting or the PreSens oxygen meter. Oxygen sensors were calibrated to 0% saturation (nitrogen purged seawater) and 100% saturation (fully aerated seawater) prior to each experiment. Background respiration was measured for 30 min (one slope) after each measurement and the run was discarded, if background respiration was >10 %. After each experiment the whole system excluding the oxygen sensors was disinfected with household bleach or Virkon® (Antec International Limited, Suffolk, United Kingdom) and rinsed with freshwater afterwards.

##### 4.3.2.3. Measurement protocol

Respiration measurements of F0 juveniles were done on approx. 5 (119 – 165 dph) and 15 months (454 – 495 dph) old juveniles. F1 juvenile respiration measurements were conducted from 2900 to 3900 dd (137-178 dph, 5 months) and 4700-5100 dd (291-318 dph, 10 months) for F1-W and F1-C, respectively. F1-C fish were older than F1-W fish at the measurement time in order to have comparable fish sizes (see Table S 4).

Juvenile sea bass were fasted for 48-72h prior to respiration measurements to minimize the effect of residual SDA (Dupont-Prinet, et al., 2010). Juveniles were randomly taken from their tank and placed individually in the respiration chambers. The whole setup was shielded from external disturbances with curtains or black foil, but the individual respiration chambers were not shielded from each other. F0 juveniles were chased until exhaustion prior to introduction to the chambers. Each experiment lasted for about 70 hours in F0 and 65 hours in F1. Oxygen partial pressure was measured 1/s and was usually kept above 80%, until start of critical oxygen concentration (*P*O_2crit_) trial (see below). Each cycle was composed of 360 s (F0) and 540 s (F1) flush time, during which oxygen concentration was restored to normoxia (until *P*O_2crit_ trial), followed by 30 s wait and 210 s (F0) and 180 s (F1) measurement time. In F0 only the measurements taken after the fish fully recovered from chasing stress were used to calculate SMR, usually after 10 hours. In F1, the first 5 hours of each experiment were not used for analysis of SMR, to account for acclimation of the fish to the respirometer and recovery from handling stress, resulting in approx. 390 and 310 cycles in F0 and F1 juveniles, respectively. Analyses were performed only on declines in oxygen concentration displaying R² > 0.85 and R² > 0.90 in F0 and F1, respectively. On the third morning, a *P*O_2crit_ trial was done on F0-old and F1 juveniles, see below. After finishing the trial or the respiration measurement for F0-young, fish were removed from the chamber. F0 juveniles were weighed and measured in BL prior to the experiment, F1 juveniles after the experiment. F0-old juveniles were identified by their PIT tag and returned to their origin tank after the experiment. F0-young juveniles and F1 juveniles were killed by a cut through the spine after the experiment.

##### 4.3.2.4. Critical oxygen concentration trial

On the third morning, oxygen concentration in the tank surrounding the chambers was continuously decreased, in F0-old by passing the water through a gas equilibration column supplied with nitrogen gas before pumping it to the tank. In F1 the decrease in oxygen concentration was done by bubbling nitrogen directly into the surrounding water bath. The decrease lasted over a period of four to six hours to determine *P*O_2crit_. When the fish lost equilibrium in the oxygen depleted chambers, they were removed from their chamber and treated as described above.

#### 4.3.3. Data handling

In F0 juveniles the metabolic rate (MR, in mg O_2_ h^-1^ kg WW^-1^ in F0) was calculated by the Aquaresp software. In F1 oxygen concentration was converted from % air saturation to nmol l^-1^ and mmol l^-1^ in larvae and juveniles, respectively (“conv_O2” function of “respirometry” package, (Birk, 2020)). MRs were calculated from the raw data with the following formulas:

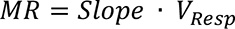

With: Slope – oxygen decline in the respiration chamber during one measurement cycle ([nmol O_2_ l^-1^ h^-1^] and [mmol O_2_ l^-1^ h^-1^] for larvae and juveniles, respectively), V_resp_ – Volume of respirometer [l].

RMR of F1 larvae was calculated as the mean MR throughout the measuring period (approx. 4h). SMR of F0 and F1 juveniles was calculated following the protocol of Chabot et al. (2016), or calculated in R with the “calcSMR” function of “fishMO2” package (Chabot, 2020). Briefly, the best SMR was chosen as described in Chabot et al. (2016). Both RMR and SMR were divided by fish mass (resulting in RMR_Raw_ and SMR_Raw_) and then corrected for allometric scaling with the following formulas:

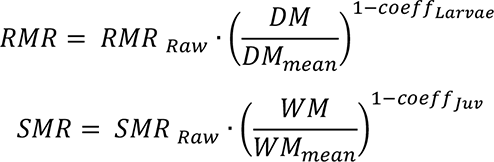

With: RMR_Raw_ and SMR_Raw_ – RMR [nmol O_2_ µg DM^-1^ h^-1^] and SMR [mmol O_2_ kg WW^-1^ h^-1^] calculated as described in the text, DM – larval dry mass [µg], WM – juvenile wet mass [kg], DM_mean_ and WM_mean_ – Mean DM and WM of all larvae and juveniles, respectively, coeff_Larvae_ and coeff_Juv_ – allometric scaling coefficient for larvae (0.89) and juveniles (0.99), respectively. The allometric scaling coefficients used were the slopes of linear regressions of MR over mass in the whole larval (F1) and juvenile (F0 and F1 together) dataset. Q_10_ was calculated with the same formula as used for SGR (see section 4.2.3).

*P*O_2crit_ was calculated with the “calcO2crit” functions of “fishMO2” package (Chabot, 2020), or according to Claireaux and Chabot (2016).

### 4.4. Statistical analysis

All statistics were performed with R (R Core Team, 2020). Significance for all statistical tests was set at *P* < 0.05. All graphs are produced with the “ggplot2” package (Wickham, 2009).

#### 4.4.1. Growth data

BL (F0 and F1 larvae and juveniles), DM (F1 larvae) and WM (F0 and F1 juveniles) data were tested for outliers (Nalimov test), normality (Shapiro-Wilk’s test, p > 0.05) and homogeneity (Levene’s test, p > 0.05). As not all assumptions for ANOVA were met, the data were fitted to linear mixed effect models (LME models, “lme” function of “nlme” package Pinheiro, et al., 2017).

Across generations, the dataset for larval BL was imbalanced, therefore it was not possible to test the effect of temperature, *P*CO_2_ condition, generation and their interaction separately, instead treatment was used as fixed variable, resulting in six groups: F0-A, F0-Δ1000, F1-C-A, F1-C-Δ1000, F1-W-A and F1-W-Δ1000. These groups, age and the interaction between group and age were included as fixed effects in the models for larval BL. The best random effect (rearing tank or no random effect) was chosen according to lowest Akaike information criteria (AIC) values. Afterwards, a backwards model selection process was used to determine the significant fixed variables and interactions.

As F1 larvae were reared in a full factorial design, temperature condition, *P*CO_2_ concentrations, age and their interactions were included as fixed effects in the model for log-transformed larval DM. Rearing tank was tested as random effect. Selection of random and variance structure, as well as model validation were done as described above for larval BL. As F0 and F1 juveniles had different temperature life histories as well as rearing conditions, their growth rates were not directly compared. Due to an imbalanced dataset in F1 juveniles, it was not possible to test the effect of temperature, *P*CO_2_ condition and their interaction separately. Instead, as for larval BL, treatment was used as fixed variable, resulting in three groups: F1-C-A, F1-C-Δ1000 and F1-W-Δ1000. For these groups, age and the interaction between group and age were included as fixed effects in the models for juvenile BL and WM. Juvenile WM was log transformed prior to analysis. Selection of random and variance structure, as well as model validation were done as described above for larval BL.

#### 4.4.2. Respirometry

RMR (F1 larvae), SMR (F0 and F1 juveniles) and *P*O_2crit_ (F0 and F1 juveniles) data were tested for outliers (Nalimov test), normality (Shapiro-Wilk’s test, p > 0.05) and homogeneity (Levene’s test, p > 0.05). As not all assumptions for ANOVA were met, the data were fitted to linear mixed effect models (LME models, “lme” function of “nlme” package, Pinheiro, et al., 2017).

As larvae were reared in a full factorial design, temperature condition, *P*CO_2_ concentrations and their interactions were included as fixed effects in the model. Respiration chamber, date of the experiment and the origin tank were tested as random effects. Additionally, in case of heterogeneity of data, variance structures were included in the random part of the model. The best random and variance structures were chosen according to lowest Akaike information criteria (AIC) values. Afterwards a backwards model selection process was used to determine the significant fixed variables and interactions.

Due to an imbalanced dataset for juvenile respirometry, it was not possible to test the effect of temperature, *P*CO_2_ condition, generation, age and their interaction separately, instead treatment was used as fixed variable, resulting in seven groups for SMR: F0-A-young, F0-Δ1000-young, F0-A-old, F0-Δ1000-old, F1-C-A, F0-C-Δ1000 and F1-W-Δ1000 and five groups for *P*O_2crit_: F0-A-old, F0-Δ1000-old, F1-C-A, F0-C-Δ1000 and F1-W-Δ1000. Selection of random and variance structure, as well as model validation were done as described above for RMR in larvae.

## 5. Results

### 5.1. Growth

Neither temperature nor *P*CO_2_ treatment had a significant effect on larval size at mouth opening stage in F1 larvae (Figure 2A and D, Table 4). During the following larval development, higher temperatures significantly increased growth if larvae were compared at the same age (dph): F1-C larvae were smaller than F0 and F1-W larvae at higher temperature (Figure 3A and B, Table 4). SGR ranged from 7.85 to 9.75 % day^-1^ for larval DM and 11.67 to 14.76 % day^-1^ for larval BL (Table 3). The higher growth rates in F1-W larvae resulted in Q_10_ of 1.67-2.12 and 1.81-2.35 for DM and BL (Table 3). *P*CO_2_ had no effect on growth of F0 and F1-C larvae, but reduced growth significantly in F1-W larvae (Table 4). Due to the longer larval duration in colder temperatures (900 dd equals 45 dph at 20°C and 60 dph at 15°C), F1-C larvae were of comparable size to F1-W-A and F0 larvae at metamorphosis. In contrast, F1-W-Δ1000 larvae were significantly smaller at metamorphosis than any other group of larvae (Figure 2B and E, Table 4).

**Figure 2.**
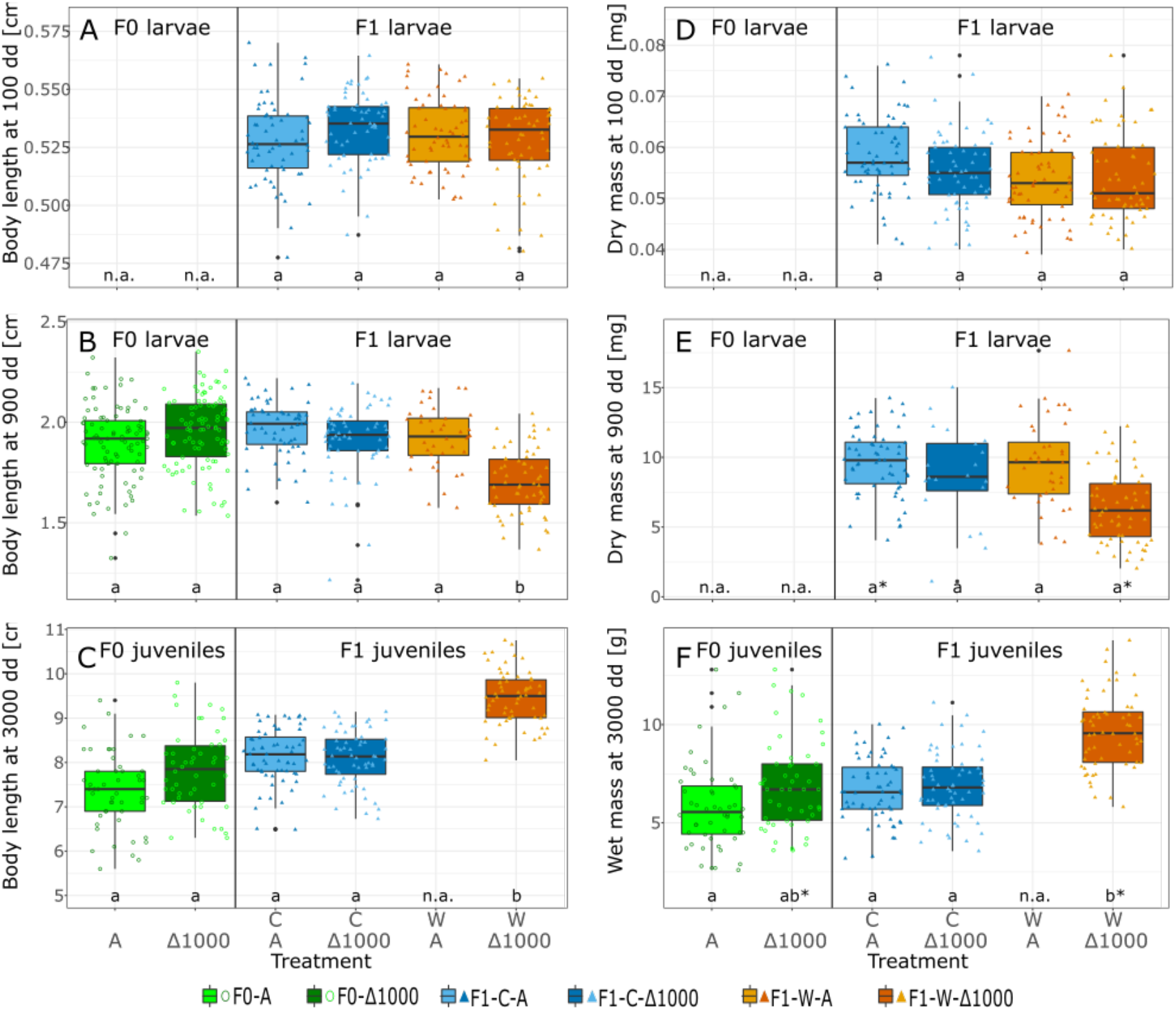
Body length and mass of European sea bass at approx. 100 dd (mouth opening, A and D, 7 dph), 900 dd (metamorphosis, B and E) and 3000 dd (C and F) in F0 and F1 fish, overlying dots are the individual data points of each treatment, different letters indicate significant differences [linear mixed effects (LME), *P*<0.05], asterisks indicate statistical trends [LME, *P*<0.1], A – Ambient *P*CO_2_, Δ1000 – ambient + 1000 µatm CO_2_, C – cold life condition, W – warm life condition, n.a. – treatment was not available or not measured at this state, n=40-90.

**Figure 3.**
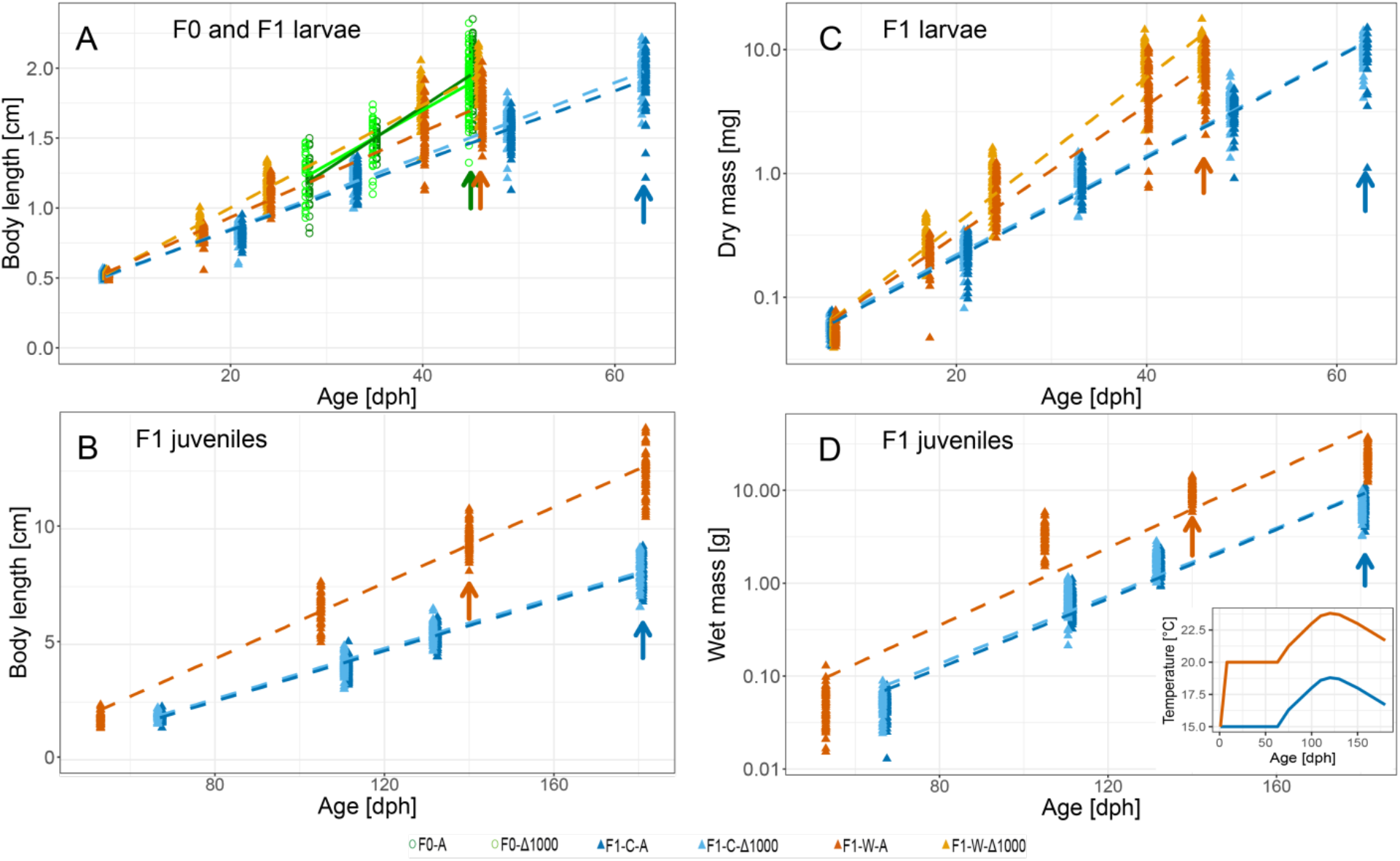
Growth of F0 and F1 larvae (A, C) and F1 juveniles (B, D) of European sea bass with linear regression lines. Shown are individual data points of body length (A, B), larval dry mass (C) and juvenile wet mass (D). F1-C larvae grew significantly slower than F1-W and F0 larvae (A, C). F1-W-Δ1000 larvae grew significantly slower than F1-W-A (A, C) and F0 larvae (A). F1-W-Δ1000 juveniles grew significantly faster than F1-C juveniles (B, D). No differences were observed between *P*CO_2_ treatments in F0 larvae (A) and F1-C larvae (A, C) and F1-C juveniles (B, D), respectively. All data were tested with LME models, F- and p-Values are summarized in Table 4. Arrows indicate the data points at metamorphosis (900dd, A, C) and at 3000 dd (B, D), data of different *P*CO_2_ conditions of the same age are slightly moved for better visibility. Insert in D shows the temperature history of F1 larvae and juveniles. A – Ambient *P*CO_2_, Δ1000 – ambient + 1000 µatm CO_2_, C – cold life condition, W – warm life condition.

**Table 3.** Specific growth rates (SGR) and their respective Q_10_ of larval and juvenile mass and body length of European sea bass. SGR [% day^-1^]and Q_10_ [-] are given for 0.05, 0.5 and 0.95 quantile of the cohort. Means ± s.e. over all replicate tanks per condition. A – Ambient *P*CO_2_, Δ1000 – ambient + 1000 µatm CO_2_, L – Larvae, J – Juveniles, C – cold life condition, W – warm life condition, n.a. – treatment was not available or not measured at this state

| Treatment | n | 0.05<br>Quantile | 0.5<br>Quantile | 0.95<sup>1011</sup><br>Quantile |
| --- | --- | --- | --- | --- |
| *Larval dry mass* |  |  |  |  |
| SGR F1 C A | 3 | 9.19±0.37 | 9.57±0.23 | 9.63±0.25 |
| SGR F1 C $\Delta 1000$ | 1 | 7.85 | 9.26 | 9.75 |
| SGR F1 W A | 2 | 12.92±0.38 | 14.25±0.11 | 14.76±0.08 |
| SGR F1 W $\Delta 1000$ | 3 | 11.67±0.29 | 12.92±0.26 | 13.73±0.20 |
| $Q_{10}$ A | | 1.84 | 1.96 | 2.12 |
| $Q_{10}$ $\Delta 1000$ | | 1.81 | 1.67 | 1.80 |
| *Larval body length* |  |  |  |  |
| SGR F1 C A | 3 | 2.27±0.08 | 2.41±0.04 | 2.40±0.06 |
| SGR F1 C $\Delta 1000$ | 3 | 2.14±0.07 | 2.33±0.02 | 2.43±0.03 |
| SGR F1 W A | 2 | 3.09±0.15 | 3.37±0.06 | 3.50±0.02 |
| SGR F1 W $\Delta 1000$ | 3 | 2.88±0.01 | 3.01±0.06 | 3.26±0.04 |
| $Q_{10}$ A | | 1.98 | 2.22 | 2.35 |
| $Q_{10}$ $\Delta 1000$ | | 1.81 | 1.95 | 1.98 |
| *Juvenile wet mass* |  |  |  |  |
| SGR F1 C A | 2 | 3.07±0.09 | 2.94±0.03 | 3.04±0.04 |
| SGR F1 C $\Delta 1000$ | 2 | 2.96±0.05 | 2.88±0.01 | 2.92±0.04 |
| SGR F1 W A |  | n.a. | n.a. | n.a. |
| SGR F1 W $\Delta 1000$ | 2 | 5.16±0.02 | 4.93±0.11 | 4.82±0.13 |
| $Q_{10}$ A | | n.a. | n.a. | n.a. |
| $Q_{10}$ $\Delta 1000$ | | 2.72 | 2.63 | 2.41 |
| *Juvenile body length* |  |  |  |  |
| SGR F1 C A | 2 | 0.91±0.02 | 0.87±0.00 | 0.87±0.00 |
| SGR F1 C $\Delta 1000$ | 2 | 0.85±0.02 | 0.84±0.00 | 0.86±0.01 |
| SGR F1 W A |  | n.a. | n.a. | n.a. |
| SGR F1 W $\Delta 1000$ | 2 | 1.55±0.01 | 1.49±0.02 | 1.46±0.06 |
| $Q_{10}$ A | | n.a. | n.a. | n.a. |
| $Q_{10}$ $\Delta 1000$ | | 2.52 | 2.45 | 2.31 |

**Table 4.** F- and p-values of fixed effects from the linear mixed models on growth and metabolic rates of F0 and F1 larval and juvenile European sea bass. n.a. – treatment was not available or not measured at this state

|  | OAW Treatment |  | PCO<sub>2</sub> Treatment |  | Temperature |  | PCO<sub>2</sub>:Temperature |  |
| --- | --- | --- | --- | --- | --- | --- | --- | --- |
|  | F-value | p-Value | F-value | p-Value | F-value | p-Value | F-value | p-Value |
| ***Larval growth*** |  |  |  |  |  |  |  |  |
| *Dry mass* |  |  |  |  |  |  |  |  |
| at mouth opening | n.a. | n.a. | 1.18 | 0.3 | 4.49 | 0.06 | 2.13 | 0.18 |
| at metamorphosis | n.a. | n.a. | 11.69 | 0.01 | 6.37 | 0.05 | 2.73 | 0.16 |
| over time | n.a. | n.a. | 17.27 | 0.0032 | 2.61 | 0.1447 | 8.01 | 0.0221 |
| *Body length* |  |  |  |  |  |  |  |  |
| at mouth opening | n.a. | n.a. | 0.28 | 0.61 | 0.25 | 0.63 | 2.54 | 0.15 |
| at metamorphosis | n.a. | n.a. | 4.70 | 0.05 | 13.10 | 0.0012 | 9.64 | 0.004 |
| over time | n.a. | n.a. | 0.07 | 0.7984 | 1009.7 | <0.0001 | 16.69 | 0.0003 |
| ***Juvenile Growth*** |  |  |  |  |  |  |  |  |
| *Wet mass* |  |  |  |  |  |  |  |  |
| at 3000 dd | 16.41 | 0.0222 | n.a. | n.a. | n.a. | n.a. | n.a. | n.a. |
| over time | 351.15 | <0.0001 | n.a. | n.a. | n.a. | n.a. | n.a. | n.a. |
| *Body length* |  |  |  |  |  |  |  |  |
| at 3000 dd | 46.93 | 0.0049 | n.a. | n.a. | n.a. | n.a. | n.a. | n.a. |
| over time | 1111.59 | <0.0001 | n.a. | n.a. | n.a. | n.a. | n.a. | n.a. |
| ***Metabolic rates*** |  |  |  |  |  |  |  |  |
| RMR | n.a. | n.a. | 0.55 | 0.46 | 45.56 | <0.0001 | 2.01 | 0.16 |
| SMR | 62.55 | <0.0001 | n.a. | n.a. | n.a. | n.a. | n.a. | n.a. |
| PO<sub>2crit</sub> | 3.08 | 0.0465 | n.a. | n.a. | n.a. | n.a. | n.a. | n.a. |

In juveniles, the overall positive effect of temperature on growth persisted, with F1-W juveniles displaying significantly higher growth rates than F1-C juveniles (Figure 3C and D, Table 4). SGR ranged from 2.88-5.16 % day^-1^ for juvenile WM and 0.84-1.55 % day^-1^ for juvenile BL, the higher growth rates resulted in Q_10_ of 2.41-2.72 and 2.31-2.52 for WM and BL, respectively, in F1-Δ1000 juveniles. If compared at the age of 3000 dd (165, 140 and 181 dph for F0, F1-W and F1-C juveniles, respectively), the difference in size was inverted compared to metamorphosis, F1-W-Δ1000 juveniles were now significantly larger than any other group (Figure 2C and F, Table 4). *P*CO_2_ did not have any significant effect on growth of F0 or F1-C juveniles. The effect of *P*CO_2_on F1-W juveniles was not determined due to the missing F1-W-A treatment.

### 5.2. Metabolic rates

Metabolic rate estimations were done on larvae with mean size ranging from approx. 1.5 to 3.0 mg DM and 11.5 to 14 mm BL with no significant differences in size between treatments (BL and DM, Table S 3). For juveniles, mean size ranged from approx. 3 to 62 g WM and 9 to 20 cm BL (Table S 4), with no significant differences in size (BL and WM) or condition factor between acidification treatments of the same age and generation (ANOVA, *P*>0.05 for F0-old; LME, *P*>0.05 for F0-young and F1-C) nor between F1-C and F1-W (LME, *P*>0.05). The positive effect of temperature on growth was mirrored in larval RMR in F1: RMR was significantly lower in F1-C compared to F1-W (Figure 4A, Table 4). But in contrast to growth no effect of *P*CO_2_ treatment or an interaction of temperature and *P*CO_2_ treatment on larval RMR was observed. A Q_10_ of 2.24 and 2.51 was calculated for larval RMR for F1-A and F1-Δ1000 larvae, respectively. Similarly, juvenile SMR was significantly lower in F1-C compared to F1-W juveniles (Figure 4B, Table 4), with Q_10_ of 1.61 for F1-Δ1000 juveniles. The comparison between the two generations showed that the SMR in the F0 juveniles did not change significantly between 5- and 15-months old juveniles, but F0-SMR estimates were significantly lower than those in F1 juveniles (Table 4). Comparable to larval RMR, there was no significant effect of *P*CO_2_ in juvenile SMR at each thermal treatment. Although the LME model states a significant effect of treatment on the critical oxygen concentration *P*O_2crit_ (Figure 4C, Table 4), posthoc tests revealed only a significant difference between F0-Δ1000 and F1-C-A (*P*<0.04), all other groups were not significantly different from each other.

**Figure 4.**
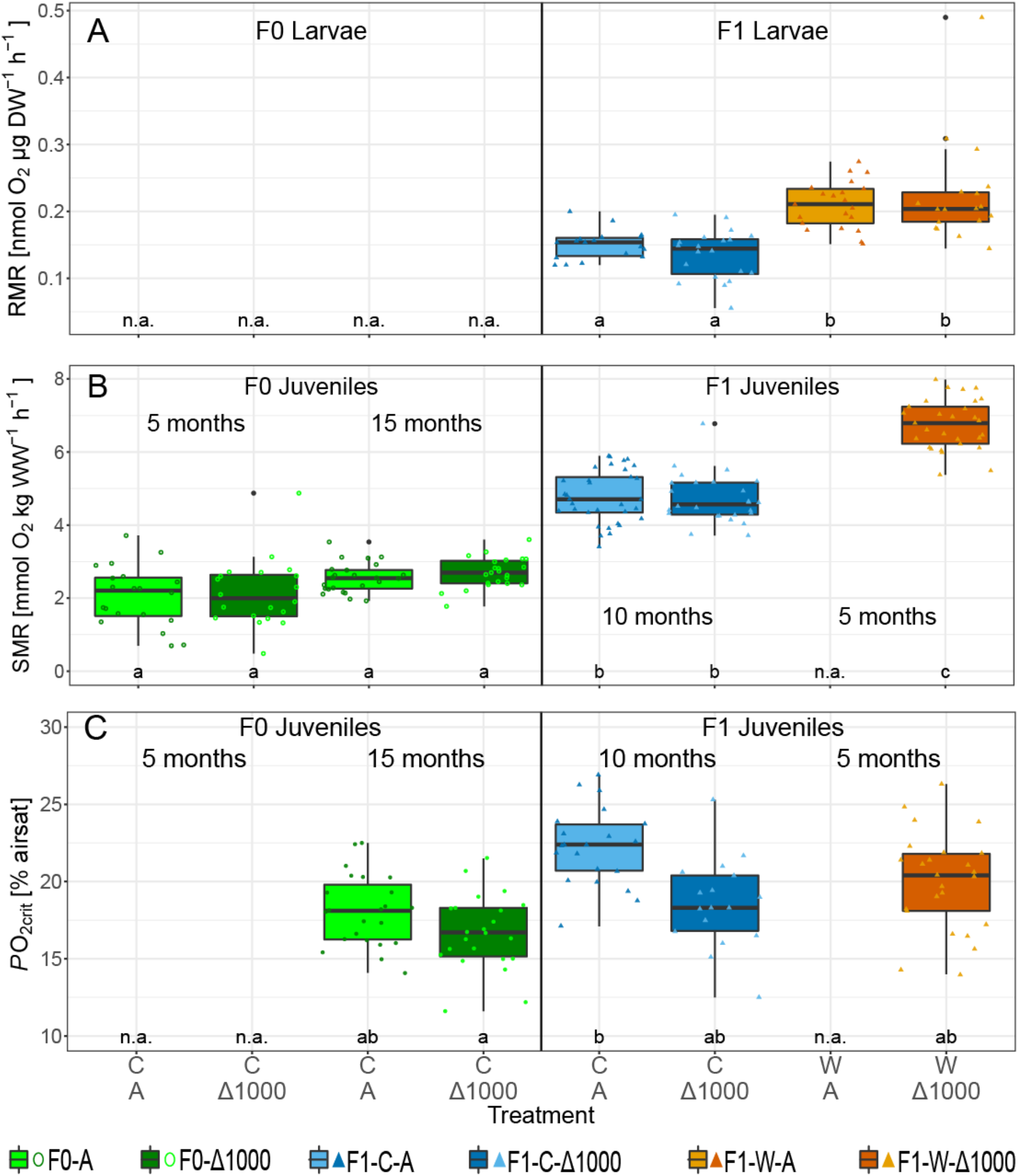
Routine (RMR, A) and standard metabolic rates (SMR, B) and critical oxygen concentration (*P*O_2crit_, C) of F0 and F1 larvae and juveniles, overlying dots are the individual data points of each treatment. Metabolic rates are corrected with allometric scaling factors (0.89 and 0.98 for larvae and juveniles, respectively). Different letters indicate significant differences [linear mixed effects (LME), *P*<0.05]; data of 15 months old F0 juveniles taken from Crespel et al. (2019). A – Ambient *P*CO_2_, Δ1000 – ambient + 1000 µatm CO_2_, C – cold life condition, W – warm life condition, n.a. – treatment was not available or not measured at this state, n=20-35.

## 6. Discussion

Long-term experiments exploring the potential of fish to adapt to OAW are still scarce, especially in larger, temperate species with long generation times. In this transgenerational experiment, we observed that OW as single driver increased growth rates and RMR in the warm F1 larval sea bass, but due to the decreased larval phase duration at warmer temperatures, F1-C-A and F1-W-A larvae had similar size at metamorphosis. OA as single driver had no effects on F1 larval and juvenile growth nor on metabolism at ambient (cold) temperature. Under OAW, F1-W-Δ1000 larvae were significantly smaller at metamorphosis than any other group, while maintaining similar RMR as F1-W-A larvae. As they grew into juveniles, F1-W-Δ1000 fish were bigger than F1-C fish at 3000 dd and had the highest SMR. Unfortunately, the F1-W-A group could not be kept until juvenile phase. Although F0 and F1-W larvae were both raised at increased temperatures, we observed that the detrimental effects of OAW occurred only in F1-W-Δ1000 and not in F0-Δ1000. We also observed that juvenile SMR was lower in F0 than in F1-C and F1-W, with no effect of OA. Juvenile *P*O_2crit_ was not affected by OA, OW or OAW in both generations.

### 6.1. Effects of OW on European sea bass growth and metabolism

F1-C larvae were reared at 15°C, reflecting ambient temperature towards middle to end of the spawning season in the Bay of Brest. We applied a warming scenario of + 5°C on F1-W larvae, which reflects typical rearing temperatures in aquaculture, as well as natural temperatures towards middle to end of the spawning season in the Mediterranean (Ayala, et al., 2003). This thermal treatment (20°C) is well below the upper thermal limits for seabass larvae from the Bay of Brest (27°C, Moyano, et al., 2017). OW as a single driver at ambient *P*CO_2_ significantly increased growth rates and decreased the time to reach metamorphosis in F1-W-A larvae in comparison to F1-C-A larvae. Due to the longer larval phase duration, size at metamorphosis was comparable between F1-C-A larvae and F1-W-A larvae. Faster growth at higher temperatures and similar size at metamorphosis despite different temperatures has been shown in other studies for sea bass from Mediterranean and Atlantic populations (Ayala, et al., 2001; 2003). OW also increased RMR in F1-W-A larvae compared to F1-C-A larvae. The increase in RMR was similar to the increase in SGR, reflected by similar Q_10_ (1.96, 2.22 and 2.24 for SGR of dry mass and body length (DM and BL, 0.5 Quantile) and RMR, respectively). This reflects the expected Q_10_ increase of 2-3 for biological processes and confirmed our hypothesis that OW will lead to increased growth and RMR in larval sea bass of this particular population. We did not determine the effects of OW as a single driver on growth and metabolism in F1 juveniles, due to the absence of F1-W-A.

### 6.2. Effects of OA on European sea bass growth and metabolism

OA as single driver at ambient temperatures did neither affect growth and metabolism (RMR; SMR), nor *P*O_2crit_ in F1 European sea bass larvae or juveniles. In the wild, sea bass eggs are spawned in stable open ocean conditions and larvae develop during the drifting towards the coast, therefore larvae were thought to be less resilient to OA than juveniles and adults. This has already been proven not to be the case for sea bass in scenarios up to SSP5-8.5 and similar (Pope, et al., 2014; F0 in Crespel, et al., 2017) and was further confirmed by this study, as larval growth and RMR were not affected by OA alone. As juvenile sea bass inhabit coastal areas and have been shown to be tolerant to a broad range of environmental factors, including temperature and salinity (Dalla Via, et al., 1998; Claireaux & Lagardère, 1999), their tolerance to OA was expected and could be confirmed in this study – no effects of OA on growth, SMR and *P*O_2crit_ were observed. Our study also supports the hypothesis of Montgomery et al. (2019) that an observed 20% decrease in *P*O_2crit_ under acute increase of *P*CO_2_ in European sea bass will vanish after long-term acclimation to OA.

### 6.3. Combined effects of OA and OW on European sea bass growth and metabolism

However, the combined effects of OA and OW (OAW) changed the picture for larval resilience. While growth rates increased sufficiently in F1-W-A to reach the same size at metamorphosis than F1-C-A, F1-W-Δ1000 larvae were significantly smaller at metamorphosis than larvae from any other treatment, but maintained RMR as high as F1-W-A larvae. Q_10_ values revealed that temperature had a stronger effect on metabolic rate than on growth under OA: 1.67 and 1.95 for SGR of DM and BL (0.5 Quantile) and 2.51 for RMR, respectively. This suggests that F1-W-Δ1000 larvae either allocated the energy differently, such as using more energy for movement or different regulatory processes, or that their energy production and oxygen usage was not as efficient as in the other groups. Although it is possible that the higher RMRs are due to higher activity of the F1-W-Δ1000 larvae during the measurements, larvae were regularly observed during the trials and the inter-individual variability in movement did not seem related to treatment. Therefore it seems more plausible that larvae under OAW needed energy for different regulatory processes, probably combined with decreased energy production efficiency. In this sense, we already found that OAW decreased the efficiency of complex II (CII) of the electron transport system (ETS) in cardiac mitochondria of juvenile sea bass in the W-Δ1000 treatment under acute temperature change (Howald, et al., 2019). Inhibition of CII by OA was also found in other studies on mammals and fish (Simpson, 1967; Wanders, et al., 1983; Strobel, et al., 2013). In Atlantic cod embryos, reduced activity of complex I (CI) of the ETS resulted in reduced mitochondrial phosphorylation capacity and subsequently in reduced oxygen consumption rates, while energy requirements were simultaneously increased (Dahlke, et al., 2016). Although CII was only affected in juvenile sea bass under acute temperature change, it is probable that larvae are more vulnerable than juveniles (Dahlke, et al., 2020): similar to embryos (Leo, et al., 2018), they are less developed while at the same time investing all available energy into growth without reserving excess capacity for environmental regulation and are therefore already affected at their acclimation temperature if OA and OW are combined. This inability to cope with OAW has not been observed in European sea bass larvae before, contrastingly in former studies growth of larval European sea bass has been shown to be resilient to OA even at rearing temperature of 19°C (Pope, et al., 2014; F0 larvae in Crespel, et al., 2017). Potential explanations why these differences first occurred in F1 are likely related to transgenerational effects, as well as effects due to different rearing protocols, which are both addressed below (section 6.4).

In contrast to larvae, F1-W-Δ1000 juveniles displayed a greater thermal plasticity and grew significantly faster than F1-C juveniles, resulting in larger fish at 3000 dd in the F1-W-Δ1000 than in F1-C-A and F1-C-Δ1000. High growth rates were supported by high SMR, which were also highest in F1-W-Δ1000 juveniles in comparison to F1-C-A and F1-C-Δ1000. As we did not incubate the F1-W-A treatment to juvenile phase, it is unclear whether the detrimental effects of OAW on growth and metabolism in larval European sea bass would have persisted into the juvenile phase. The increased growth rates and bigger size at 3000 dd in F1-W-Δ1000 juveniles in comparison to F1-C-A and F1-C-Δ1000 juveniles might either indicate that OA did not affect growth in juveniles or that growth under OW was so much accelerated in juveniles that F1-W-Δ1000 fish were able to catch up and grow to bigger sizes than F1-C fish masking the negative effects of OAW. The latter suggestion is supported by the findings in SMR and by the Q_10_ of SMR and SGR: in F1-Δ1000 juveniles, SMR was less affected by temperature (Q_10_ 1.61) than SGR (Q_10_ 2.63 and 2.45 for SGR of WM and BL (0.5 Quantile)). Q_10_ for SGR and SMR are well in the range found in other studies on European sea bass from the Atlantic (Q_10_ for SGR of WM ∼ 2.4 (15-20°C, calculated from Gourtay, et al., 2018) and Q_10_ for SMR 2.09 (14-22°C, Montgomery, et al., 2021)) and from the Western Mediterranean populations (Q_10_ for SGR of WM and RMR of 2.40 and 1.70, respectively, 13-25°C, calculated from Person-Le Ruyet, et al., 2004). The authors of the latter study explained the different Q_10_ of RMR and SGR with increased growth rates due to increased feed intake. As the fish in our study were fed *ad libitum*, they were able to increase food intake to support high growth rates, too. The better capacity of juveniles to cope with and even profit from higher temperatures even under OAW in comparison to larvae is probably due to the reproduction biology of European sea bass, as well as to the generally higher capacity for acid-base regulation in juveniles in comparison to larvae. Larvae are developing during spring in the open ocean resulting in stable and relatively cold temperatures (8-13°C for Atlantic specimen, Jennings & Pawson, 1992), with optimal larval growth temperatures of 15-17°C (Mediterranean specimens, Koumoundouros, et al., 2001; Ayala, et al., 2003). Juveniles on the other hand live in shallow coastal areas, resulting in higher temperatures during summer but also higher daily and seasonal variation (6-18°C for Atlantic specimens, Russel, et al., 1996) with optimal growth temperatures of 22-28°C (Mediterranean specimens, Lanari, et al., 2002; Person-Le Ruyet, et al., 2004). Consequently, in terms of growth and metabolism juvenile sea bass at the northern distribution range might benefit from higher temperatures, as already found in other studies (Howald, et al., 2019; Montgomery, et al., 2021) and do not seem to be severely affected by OA.

### 6.4. Transgenerational effects under OA on European sea bass growth and metabolism

In addition to the single and combined stressors OA, OW and OAW, we also studied the transgenerational effects on the ability of sea bass larvae and juveniles to cope with upcoming conditions. This study is to our knowledge the first one to examine the transgenerational effects of ocean acidification (OA) on European sea bass or other long-lived teleost with parents being reared from early larval stage under OA conditions as well as their offspring. Interestingly the detrimental effect of OAW on larval growth was only observed in F1 and not in F0 larvae of European sea bass, despite their respective parental generation’s identical thermal history, and thus appears to be an OA effect. This can be explained by several reasons: First, the provisioning of necessary resources when parents have already encountered the same conditions as the future offspring, e.g. via egg size and composition (Munday, 2014) could explain the observed trend in F1-W-Δ1000 larvae. Parental effects can lead in different directions and can last throughout larval phase: parental effects influence growth in stickleback under OW and OA (Shama, et al., 2014; Schade, et al., 2014) and explained differences in embryo mortality and hatching success in Atlantic cod under OW (Dahlke, et al., 2016). In our study, we did not directly quantify parental effects, but the size of F1 larvae at mouth opening, a landmark until which the larvae depend on yolk sac reserves, did not differ across treatments. Thus parental provisioning does not seem to explain differences in larval growth rates. Second, the incubation protocol differed between F0 and F1. While F0 larvae were first incubated under OA conditions at 2 dph, F1 sea bass were constantly reared under OA conditions from fertilization onwards, although warming was also applied from 2 dph onwards. It is possible that effects of OA during embryogenesis shaped the reaction of F1 larvae to OAW, e.g. via epigenetic signaling. As reviewed by Dahlke et al. (2020), it seems that spawning adults and embryos are the most vulnerable life stages in fish, possessing the lowest tolerance to OW, e.g. Atlantic cod embryos exposed to OAW showed reduced hatching success and oxygen consumption rates (Dahlke, et al., 2016) and OA decreased the Q_10_ of RMR in Atlantic silverside embryos (Schwemmer, et al., 2020). To summarize, the different reaction of F0 and F1 larvae to OAW could be due to parental effects or effects during embryogenesis and more research is necessary to determine the underlying mechanisms.

As the different temperature life histories did not allow a direct comparison of growth rates between F0 and F1 juveniles, we compared size at the age of approx. 3000 dd (165, 140 and 181 dph for F0, F1-W and F1-C juveniles). Due to their high growth rates during juvenile phase, F1-W-Δ1000 fish were largest at 3000 dd, while F0 and F1-C fish were smaller (WM and BL) but similar to each other. This matched the result of SMR, which was not affected by OA and was higher in F1-W compared to F1-C fish. Surprisingly, SMR was also higher in F1-C compared to F0 fish. This difference might be explained by the different temperature life histories. While F0 fish had been raised at warmer temperatures and were acclimated to colder temperatures afterwards, F1-C fish had been reared at 15°C throughout their life, except for summer months, when temperatures reached up to 19°C. No detrimental effects on juvenile growth rate under OA were visible in the second generation of sea bass reared under OA conditions, reflected by similar SMR and SGR between A and Δ1000 condition. Due to the missing F1-W-A treatment, we cannot state if the detrimental effects of OAW observed in F1 larvae persisted to juvenile phase.

### 6.5. Ecological perspective

Larvae are not fully developed compared to later stages and are exposed to higher predation and starvation risks, as such they had been thought to be more vulnerable to environmental stressors such as OAW (as reviewed in Houde, 2019). In this context, OAW could impact larval survival and recruitment success via different mechanisms. If OAW leads to faster growth rates and increased metabolic rates (as seen in this study between F1-W and F1-C larvae), larvae need more food in shorter time to support these growth rates, therefore it is essential that they match adequate prey fields (prey abundance, size and quality). In our study the larvae were fed *ad libitum* at both temperatures, supporting increased energetic demands for the high growth and metabolic rates at higher temperatures. However, in the ocean it is possible that food availability is not sufficient to support accelerated growth under OW. Bochdansky et al. (2005) showed that fish larvae with higher growth and metabolic rates died earlier, when food was limited, but profited when fed at saturation level. In sea bass larvae, high growth rates were also only supported under high food ratios, but survival was not significantly decreased, even at one eighths of saturation ratio (Zambonino Infante, et al., 1996). This might indicate that sea bass will not grow as fast as in our study under future OW scenarios if food is scarce, but might still survive to juvenile stage.

Besides food-related aspects, OAW can also have a large impact on larval behavior and dispersal, which can later influence recruitment success. Sea bass spawn in the open ocean and larvae are drifted inshore (Jennings & Pawson, 1992). As with many temperate species, their swimming behavior and its effect on dispersal has not been studied as extensively as for coral reef fish that have well developed sensory abilities (hearing, olfaction, vision) and show directional swimming early on (as reviewed in Leis, 2018; Berenshtein, et al., 2021). To the best knowledge, it seems that early seabass larvae are more dependent on currents than on their swimming performance and that they are able choose a certain depth and therefore a certain current in the preferred direction (Jennings & Pawson, 1992). When being drifted closer to the coast, sea bass larvae wait for certain cues from nursery areas, which are present from June onwards (Jennings & Pawson, 1992).

OW accelerates development of sea bass larvae and therefore possibly alters the timing and spacing of dispersal. Studies have shown species-specific responses of fish behavior to OA, OW and OAW, e.g. OW increased activity level in larval kingfish but not boldness, while OA had no effect on these behavioral traits (Laubenstein, et al., 2019). Yet, OA decreased swimming duration and orientation in larval dolphinfish (Pimentel, et al., 2014) and reversed orientation towards settlement habitat cues in barramundi (Rossi, et al., 2015). To our knowledge, larval sea bass behavior has not been measured under OAW yet. Consequently due to the altered timing of larval development and in combination with the possibility of altered behavior and impacted senses, reaching nursery areas might be challenging for sea bass larvae under OAW, especially if (1) food is not abundant and (2) cues are weaker and/or different due to greater distance and/or earlier timing. Once the larvae entered the coastal areas and metamorphose, they are exposed to a more changing environment. Although this study could confirm that juvenile sea bass are less vulnerable to OAW than larval sea bass, food availability and behavior will determine, if the observed increased growth under OAW in F1 will occur in the wild, too. In a sister study on offspring of wild caught European sea bass, OAW reduced digestive enzyme activity under restricted food ratios resulting in severely reduced food conversion efficiency and reduced growth rates (Cominassi, et al., 2020). Additionally, OA decreased the distance which early juvenile sea bass needed to sense food or predator cues (Porteus, et al., 2018) and juvenile sea bass behavior was altered by OW resulting in decreased latency of escape response and mirror responsiveness (Manciocco, et al., 2015). Consequently, although faster larval (OW) and juvenile growth (OAW) as well as earlier metamorphosis (OW, OAW) is generally beneficial for larvae and early juveniles, many factors may modulate this effect and whether it will translate into higher larval survival, recruitment and increased growth rates in the wild. Further research should determine the effects of limited food under OAW on larval and juvenile growth and behaviour.

As the hypoxia tolerance of European sea bass juveniles was unaffected by OA, OW and OAW, they might cope well with upcoming hypoxia events in coastal areas. However, it is important to note here that we measured *P*O_2crit_ only at SMR and thus may have estimated *P*O_2_ effects too conservatively. Recent studies suggest that this *P*O_2crit_ at SMR might not be the most ecologically relevant estimate (see Seibel & Deutsch, 2020 and references therein): Long-term survival of individuals and their population would require that the fish are able to digest food, grow and reproduce, which would require more energy than provided by SMR. Consequently, depending on the duration and intensity of hypoxia events, individuals might be able to survive over short terms, but other fitness related traits such as growth might be affected in the long-term.

## 7. Conclusion

We confirmed our hypotheses that OW increases growth and metabolism in the European sea bass, and that larvae as well as juveniles are resilient to OA if it occurs as a single stressor. Yet, we could also confirm that OAW had detrimental effects on larval growth. Our results together with other findings on larval fish and European sea bass suggest that it is possible that under OAW fewer individuals will reach metamorphosis, e.g. due to limited food to support high growth rates, different dispersal to nursery areas by altered developmental timing, changed behavior or affected olfactory senses. However, those individuals that reach the juvenile phase might benefit from higher temperatures, due to increased performance.

## 2. List of abbreviations

Δ1000: Acidification condition (ambient *P*CO_2_ + 1000 µatm)
A: Ambient *P*CO_2_ condition
BL: Body length
C: Cold life conditioned group
CI: Complex I of the ETS (NADH dehydrogenase)
CII: Complex II of the ETS (succinate dehydrogenase)
dph: Days post hatch
dd: Degree days
DM: Dry mass
ETS: Electron transport system
IPCC: Intergovernmental Panel on Climate Change
MS-222: Tricaine methanesulfonate
OA: Ocean acidification
OAW: Ocean acidification and warming
OW: Ocean warming
*P*CO_2_: Partial pressure of CO_2_
*P*O_2_: Partial pressure of O_2_
*P*O_2crit_: Critical oxygen concentration
RCP: Representative concentration pathway
RMR: Routine metabolic rate
SDA: Specific dynamic action
SMR: Standard metabolic rate
W: Warm life conditioned group
WM: Wet mass

## 8. Acknowledgements

We thank Luis Kuchenmüller for his support in measuring larval dry mass and Mark Suquet for his support during the reproduction period. We acknowledge the technicians and researchers from the Laboratory of Adaptation, and Nutrition of Fish for their support in animal welfare during the weekends. We want to specially thank Patrick Quazuguel and Nicolas LeBayon for their help in rearing the fish from larvae to reproduction and beyond.

## 10. Author contributions

Conceptualization: SH, AC, MM, GC, MP, FCM

Data curation: SH

Formal analysis: SH, AC, MM, FCM

Funding acquisition: MP, GC, FCM

Investigation: SH, AC, LC

Methodology: SH, MM, GC, FCM

Project administration: MM, GC, MP, FCM

Resources: GC, MP, FCM

Software: SH, MM, GC

Supervision: MM, GC, MP, FCM

Validation: SH, MM, FCM

Visualization: SH

Writing - original draft: SH

Writing - review & editing: SH, MM, AC, LC, GC, MP, FCM

## 12. Data availability

Datasets of growth, metabolic rates and water conditions during rearing are available online from PANGAEA (www.pangaea.de)

## 15. Supplement

**Table S 1.** Light intensity during rearing phase of European sea bass larvae.

| Age [d] | 2 | 8 | 11 | 20 | 30 | 32 | 36 | 46 |
| --- | --- | --- | --- | --- | --- | --- | --- | --- |
| Light intensity [lux] | 0 | 0-1 | 1 | 7 | 10 | 31 | 59 | 96 |

**Table S 2.** Larval mortality in % in the different larvalrearing tanks (n=3); A – Ambient *P*CO_2_ andΔ1000 – ambient + 1000 µatm CO_2_, T – temperature, Rep 1-3 – replicate tank 1-3.

| A |  |  |  | Δ1000 |  |  |
| --- | --- | --- | --- | --- | --- | --- |
| T [°C] | Rep 1 | Rep 2 | Rep 3 | Rep 1 | Rep 2 | Rep 3 |
| 15 | 73.5 | 28.8 | 83.1 | 67.9 | 55.0 | 47.2 |
| 20 | 96.4 | 76.2 | 25.8 | 59.3 | 52.5 | 53.7 |

**Table S 3.** Biometrical data of larvae used for respiration measurements: Treatments: C – cold life condition (15°C), W – warm life condition (20°C), A – ambient *P*CO_2_, Δ1000 – ambient *P*CO_2_ + 1000 µatm, values are means ± s.e.m. Different letters indicate significant differences between groups (LME, *P*<0.05).

| Treatment | n | Dry weight<br>[mg] | Bodylength<br>[mm] |
| --- | --- | --- | --- |
| C – A | 18 | 2.87±0.51<sup>a</sup> | 13.96±0.77<sup>a</sup> |
| C – $\Delta 1000$ | 20 | 2.95±0.46<sup>a</sup> | 14.04±0.80<sup>a</sup> |
| W – A | 21 | 2.51±0.43<sup>a</sup> | 13.04±0.71<sup>a</sup> |
| W – $\Delta 1000$ | 18 | 1.70±0.53<sup>a</sup> | 11.63±0.85<sup>a</sup> |

**Table S 4.** Biometrical data of juveniles used for respiration measurements: Treatments: C – cold life condition (up to 18°C), W – warm life condition (up to 23°C), A – ambient *P*CO_2_, Δ1000 – ambient *P*CO_2_ + 1000 µatm, values are means ± s.e.m. Different letters indicate significant differences between groups (LME, P<0.05).

| Generation | Treatment | Age [m] | n | Fish mass [g] | Forklength [mm] | Condition factor [-] |
| --- | --- | --- | --- | --- | --- | --- |
| F0 | C – A | 6 | 20 | 5.06±0.24<sup>a</sup> | - | - |
| F0 | C – $\Delta 1000$ | 6 | 20 | 5.85±0.27<sup>a</sup> | - | - |
| F0 | C – A | 18 | 24 | 81.80±2.60<sup>b</sup> | 18.11±0.18<sup>b</sup> | 1.37±0.02<sup>b</sup> |
| F0 | C – $\Delta 1000$ | 18 | 24 | 81.40±3.22<sup>b</sup> | 18.25±0.22<sup>b</sup> | 1.33±0.03<sup>b</sup> |
| F1 | C – A | 10 | 33 | 15.00±0.69<sup>c</sup> | 10.86±0.14<sup>c</sup> | 1.14±0.02<sup>c</sup> |
| F1 | C – $\Delta 1000$ | 10 | 26 | 13.05±0.56<sup>c</sup> | 10.31±0.14<sup>c</sup> | 1.17±0.01<sup>c</sup> |
| F1 | W – $\Delta 1000$ | 5 | 29 | 15.73±1.01<sup>c</sup> | 11.04±0.20<sup>c</sup> | 1.12±0.02<sup>c</sup> |

**Figure S 1.**
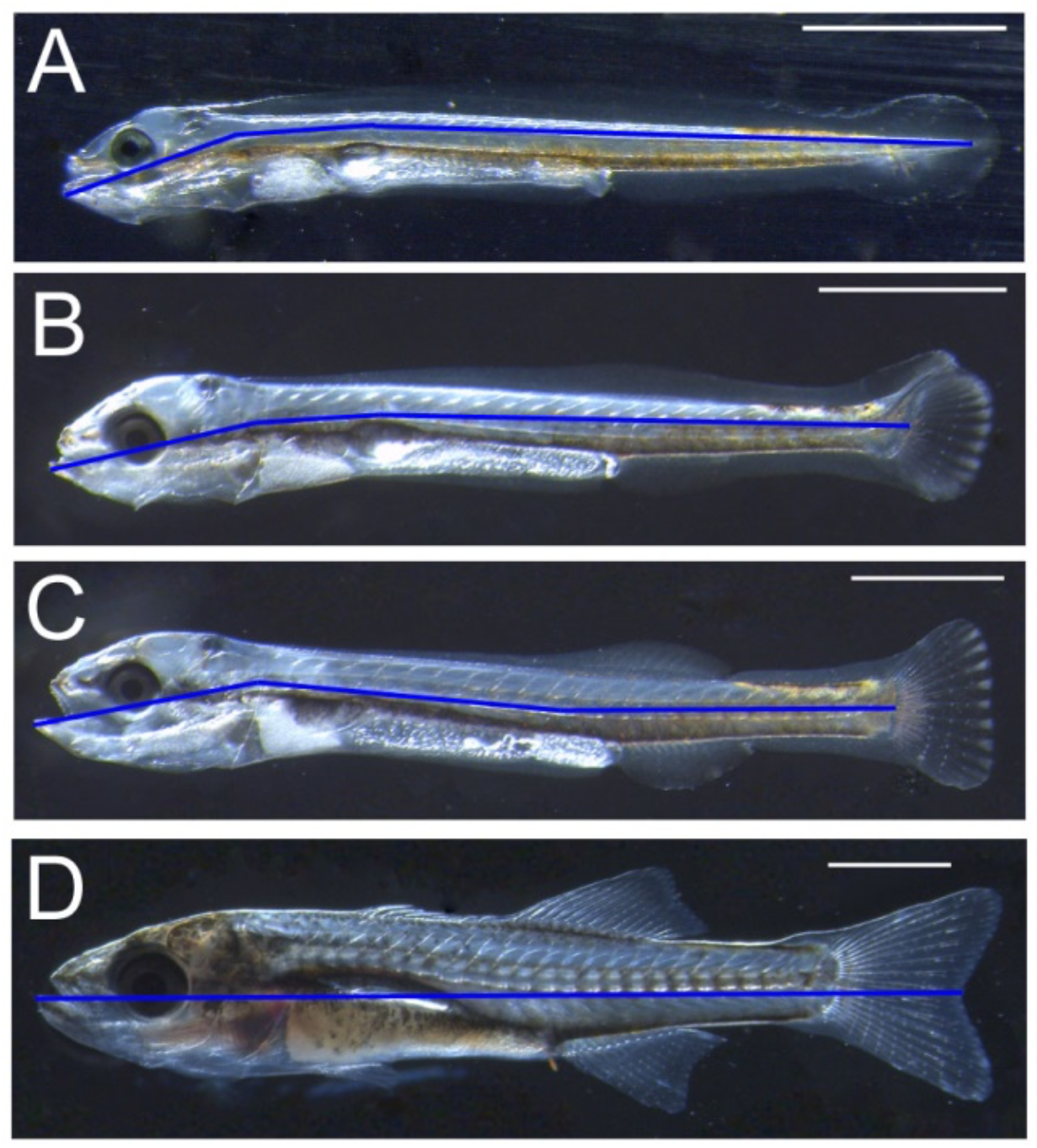
Body length measurements in larvae at different developmental stages: A – pre flexion (about 300 dd), B – flexion (about 460 dd), D-post flexion (about 460 dd) and (post)metamorphosis (about 900 dd). Until post flexion the segmented line tool in the software ImageJ (Schneider, et al., 2012) was used to measure the length of the larva, afterwards the length of the larvae was measured as a straight line, as it would be done with calipers. The lines of the measurement are marked in blue.

